# Improved alpha-beta power reduction via combined Electrical and Ultrasonic stimulation in a Parkinsonian Cortex-Basal Ganglia-Thalamus Computational Model

**DOI:** 10.1101/2021.02.03.429377

**Authors:** Thomas Tarnaud, Wout Joseph, Ruben Schoeters, Luc Martens, Emmeric Tanghe

**Affiliations:** Department of Information Technology (INTEC-WAVES/IMEC), Ghent University/IMEC, Technologypark 126, 9052 Zwijnaarde, Belgium.

**Keywords:** Ultrasonic neuromodulation, deep brain stimulation, basal ganglia, intramembrane cavitation, computational modeling

## Abstract

**Objective:** To investigate computationally the interaction of combined electrical and ultrasonic modulation of isolated neurons and of the Parkinsonian cortex-basal ganglia-thalamus loop.

**Methods:** Continuous-wave or pulsed electrical and ultrasonic neuromodulation is applied to isolated Otsuka plateau-potential generating subthalamic nucleus (STN) and Pospischil regular, fast and low-threshold spiking cortical cells in a temporally-alternating or simultaneous manner. Similar combinations of electrical/ultrasonic waveforms are applied to a Parkinsonian biophysical cortex-basal ganglia-thalamus neuronal network. Ultrasound-neuron interaction is modelled respectively for isolated neurons and the neuronal network with the NICE and SONIC implementations of the bilayer sonophore underlying mechanism. Reduction in *α—β* spectral energy is used as a proxy to express improvement in Parkinson’s disease by insonication and electrostimulation.

**Results:** Simultaneous electro-acoustic stimulation achieves a given level of neuronal activity at lower intensities compared to the separate stimulation modalities. Conversely, temporally alternating stimulation with 50 Hz electrical and ultrasound pulses is capable of eliciting 100 Hz STN firing rates. Furthermore, combination of ultrasound with hyperpolarizing currents can alter cortical cell relative spiking regimes. In the Parkinsonian neuronal network, high-frequency pulsed separated electrical and ultrasonic deep brain stimulation (DBS) reduce pathological *α* — *β* power by entraining STN-neurons. In contrast, continuous-wave ultrasound reduces pathological oscillations by silencing the STN. Compared to the separated stimulation modalities, temporally simultaneous or alternating electro-acoustic stimulation can achieve higher reductions in *α* — *β* power for the same contraints on electrical/ultrasonic intensity.

**Conclusion:** Continuous-wave and pulsed ultrasound reduce pathological oscillations by different mechanisms. Electroacoustic stimulation further improves *α*— *β* power for given safety limits and is capable of altering cortical relative spiking regimes.

**Significance:** focused ultrasound has the potential of becoming a non-invasive alternative of conventional DBS for the treatment of Parkinson’s disease. Here, we elaborate on proposed benefits of combined electro-acoustic stimulation in terms of improved dynamic range, efficiency, resolution, and neuronal selectivity.

## I. Introduction

IN the last decade, neuromodulation by ultrasound (UNMOD) has become more popular, due to its high spatial resolution (millimeter resolution in the transversal direction with a single transducer), non-invasiveness, reversibility and safety [1]–[6]. Furthermore, phased-arrays of ultrasound transducers have been used for the non-invasive ablation of brain tumours or for subthalamotomy (high intensity focused ultrasound) [7]–[10]. Consequently, similar technology could in theory be used with Low Intensity Low Frequency Ultrasound (LILFU) in order to target deep brain structures non-invasively for neuromodulation [4], [11]–[13].

For the treatment of Parkinson’s disease, the subthalamic nucleus (STN) is an important target, also used in conventional (electrical) deep brain stimulation (DBS). In electrical DBS, an electrode lead is surgically implanted in the brain and connected with an implanted pulse generator via wires that run subcutaneously. Electrical pulses will then modulate the activity of the neuronal tissue surrounding the implanted lead. Conventional DBS has proven effective for the improvement of Parkinsonian symptoms and motor scores. However, the surgery carries a risk of complications such as infection or haemmorhage [14]–[17]. Here, the potential application of transcranial LILFU for non-invasive deep brain stimulation has been considered before [4], [11]–[13], [18]. Furthermore, recent *in vivo* studies in MPTP (1-methyl-4-fenyl-1,2,3,6-tetrahydropyridine) lesioned Parkinsonian mice have demonstrated ultrasound-induced striatal dopamine normalization [19], [20], restored locomotion activity (open field test, pole test) [19]–[22], and an increase in striatal total superoxide dismutase (T-SOD) and glutathione peroxidase (GSH-PX) (neuroprotective antioxidant enzymes) [21]. Moreover, ultrasound focused at the STN or globus pallidus (GP) downregulates the Bax to Bcl-2 ratio (Bax and Bcl-2 are proapoptotic and antiapoptotic, respectively), resulting in a reduction of cleaved-caspase 3 activity in the substantia nigra [22]. Also, *in vitro,* an increased dopamine release in PC12 cells upon low-intensity continuous insonication is observed in [19]. These first studies on ultrasonic neuromodulation for Parkinson’s disease, indicate that transcranial LILFU could be a promising alternative to conventional electrical DBS or L-DOPA (L-3,4-di-hydroxy-phenylalanine) therapy. Here, UNMOD could be applied for patient selection or to optimize the choice of the deep brain target for conventional DBS.

However, the underlying mechanism of ultrasonic neuromodulation is not well understood. Several tentative mechanisms have been proposed, e.g., bilayer sonophores [23]–[25], acoustic-radiation pressure [26]–[28], mechanosensitivity of protein channels [29], [30], flexoelectricity [31], extracellular cavitation [32], and thermodynamically-based models [33]. Here, computational models will help in order to compare the predictions of a proposed mechanism with experimental observations. Similarly, computational models have been used to improve understanding of deep brain stimulation and Parkinson’s disease, i.a., DBS-induced normalization of thalamic relay [34], [35], beta-oscillatory power [36], [37] and GPi-bursting [38], importance of antidromic activation of the hyperdirect pathway [39], [40], short-term synaptic depression and axonal failure [41], mechanisms of the generation and interaction of Parkinsonian beta-oscillations [37], [42], and effects on learning and impulsivity [43], [44].

Consequently, in this study, our goal is to achieve the following two novelties. First, to investigate the capability and efficiency of ultrasonic neuromodulation to reduce beta-oscillatory power, as a proxy for Parkinsonian bradykinesia, in a computational model of the cortex-basal ganglia-thalamus neuronal (CTX-BG-TH) network. Here, we implemented a fully biophysical Hodgkin-Huxley based network of the cortex-basal ganglia-thalamus, based on earlier models of the basal ganglia [34]–[36] and of the cortex [25], [45]–[47]. In particular, a computational spiking-neuron model (as opposed to firing rate based models) of the CTX-BG-TH has been proposed in [36], with integrate-and-fire representations for the cortical cells. We included biophysical cortical Pospischil-models [45] with cortical short-term synaptic plasiticity [46]. We opted for a fully Hodgkin-Huxley network in order to have a description of ionic channels, allowing future investigations on the mechanisms of UNMOD in the neuronal network via the interaction with mechanosensitive ion channels. For ultrasonic neuromodulation, we focus in this study on the bilayer sonophore model of intramembrane cavitation [23]– [25]. We investigate the relative efficiency for the reduction of Parkinsonian beta-oscillations with pulsed or continuous ultrasound, compared to conventional electrical deep brain stimulation.

Second, we explore the potential benefits of applying ultrasonic neuromodulation combined with electrostimulation in tandem. Recently, in [48], ultrasonic neuromodulation was combined with transcranial magnetic stimulation (TMS) to investigate the effect of ultrasound on single-pulse motor evoked potentials and on paired-pulse TMS-metrics, such as short interval intracortical inhibition (SICI) and intracortical facilitation (ICF). In theory, UNMOD could potentially be combined with all electrostimulation technologies (TMS, tDCS, DBS, etc.), with the dual aim of improving understanding of these technologies and improving their therapeutic effects (safety, resolution, etc.). For the application of non-invasive transcranial deep brain stimulation, an idea is to combine ultrasound with temporal interference deep brain stimulation (TI-DBS) [12], [49]. In TI-DBS, two high-frequency (*f*_1_ = 2 kHz and *f*_2_ = *f*_1_ + 10 Hz) electrical currents are applied transcranially, resulting in maximal modulation depth by temporal interference at the targeted deep brain region, allowing non-invasive deep stimulation in mice. However, computational studies have predicted that results of TI-DBS in humans might be less favourable [50], [51]. Besides improving the placement, waveforms, and number of the TI-DBS anode-cathode pairs, another option could be to combine the benefits of TI-DBS and UNMOD by application in tandem. Here, to our best knowledge for the first time, we study the response to combined ultrasonic and electrical neuromodulation in isolated neuron models. Then, we investigate the benefits of their combined application in the neuronal CTX-BG-TH network.

## II. Methods

### A. Cortex-Basal Ganglia-Thalamus Neuronal Network

A computational neuronal network model of the cortex-basal ganglia-thalamus loop was realised in the DynaSim MATLAB^®^-toolbox for neural modeling and simulation [52] (Euler discretisation, time step Δ*t* = 10 *μ*s for the Parkinsonian network, electrical DBS and continuous-wave ultrasound, Δ*t* = 5 *μ*s for ultrasonic DBS, and Δ*t* = 2.5 *μ*s for combined ultrasound and electrostimulation: a lower temporal discretization step is used for ultrasonic or combined electroacoustic stimulation, due to higher computational stiffness). All simulations are run for 11 s, where initialisation-dependent network transient effects during the first second are removed. A schematic overview of the network topology is given in Fig. 1(a). The cortical network (CTX) is based on [25], [46], [47] and consists of twenty regular spiking (RS), five fast spiking (FS), and five low-threshold spiking (LTS) cells. Cortical cell numbers are in agreement with estimates that about 30% of cortical cells are interneurons and 50% of interneurons are fast spiking [46], [53]. Basal ganglia (BG) cells included in the network are ten indirect striatal (iStr), ten direct striatal (dStr), twenty globus pallidus external (GPe), twenty globus pallidus internal (GPi) and twenty subthalamic nucleus (STN) neurons. The basal ganglia network model is based on [34]–[36], [54]. Finally, twenty thalamic (Th) cells receive afferent input from the basal ganglia output nucleus (GPi) and are connected with the regular and fast spiking cortical cells [47].

**Fig. 1:**
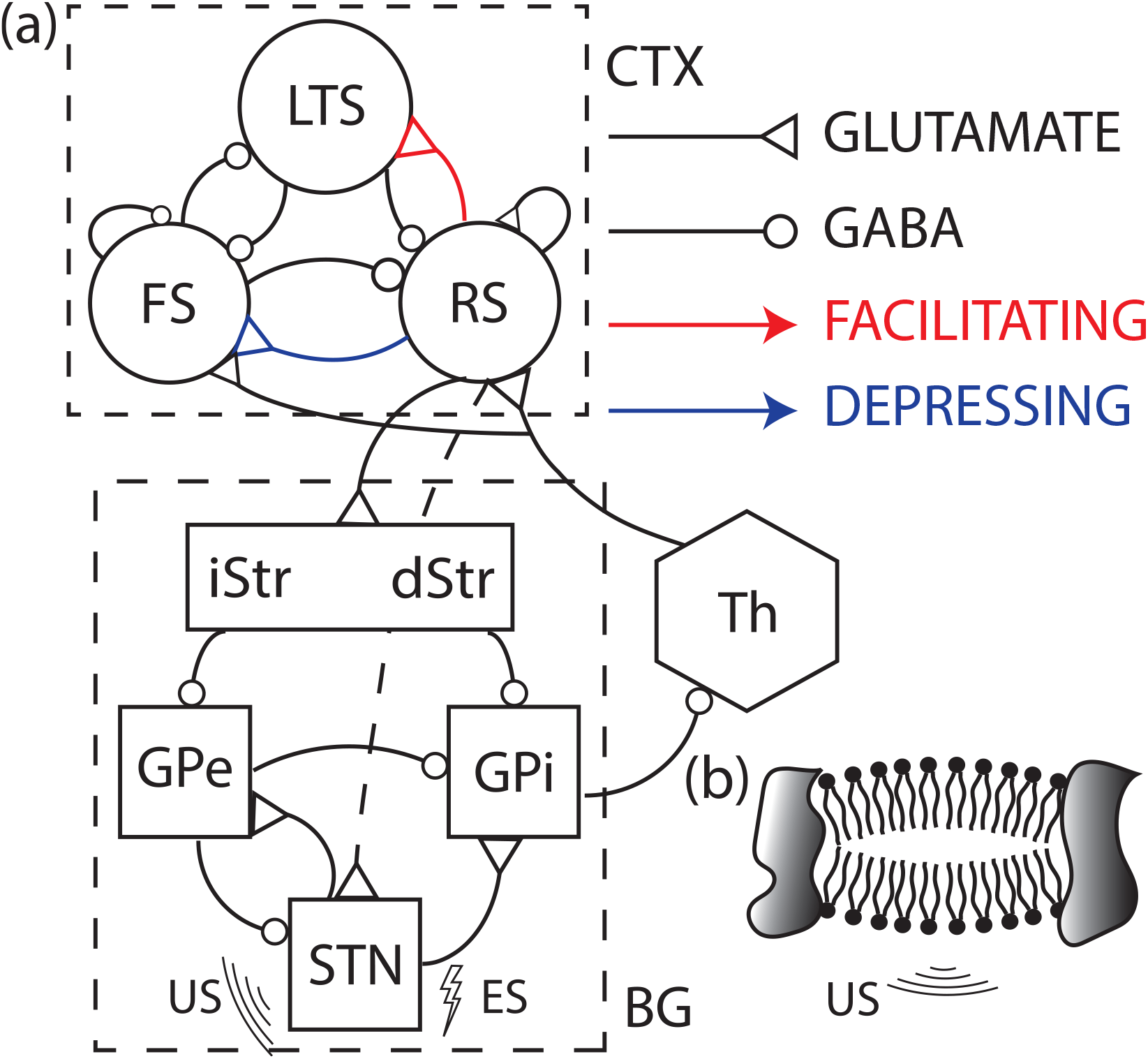
(a) Schematical overview of the cortex-basal ganglia-thalamus model. GABAergic inhibitory and glutamatergic (AMPA or NMDA) excitatory synapses are depicted with circular and triangular arrows, respectively. Red and blue arrows are used to indicate facilitating and depressing short term synaptic plasticity. The dotted RS-STN line represents the hyperdirect pathway. (b) Illustration of the bilayer sonophore interaction with ultrasound. The ultrasound-induced oscillations of the biphospholipid layer is constrained by the surrounding protein islands. **Abbreviations.** CTX: cortex, RS: regular spiking, FS: fast spiking, LTS: low threshold spiking, iStr: indirect striatum, dStr: direct striatum, GPe: globus pallidus externus, GPi: globus pallidus internus, STN: subthalamic nucleus, Th: thalamus, US: ultrasonic stimulation, ES: electrical stimulation, GABA: gamma-aminobutyric acid, AMPA: alpha-amino-3-hydroxy-5-methyl-4-isoxazolepropionic acid, NMDA: N-Methyl-d-aspartic acid.

#### 1) Network topology

Cortical network topology is based on [46], but with point neurons and a rescaled number of neurons. Synaptic connections between cortical cells are assigned by selecting source neurons at random (excluding self-connections) with network connectivity (netcon) probabilities summarized in Table Ia [46]. The number of connections between the cortex, basal ganglia and thalamus subsystems is given in Table Ib. Basal ganglia model topology is deterministic: the number of synaptic connections is summarized in Table Ic [36]. Here, as in Kumaravelu et al. (2016), 50% of globus pallidus cells chosen at random will not receive subthalamic nucleus afferents.

**TABLE I:**
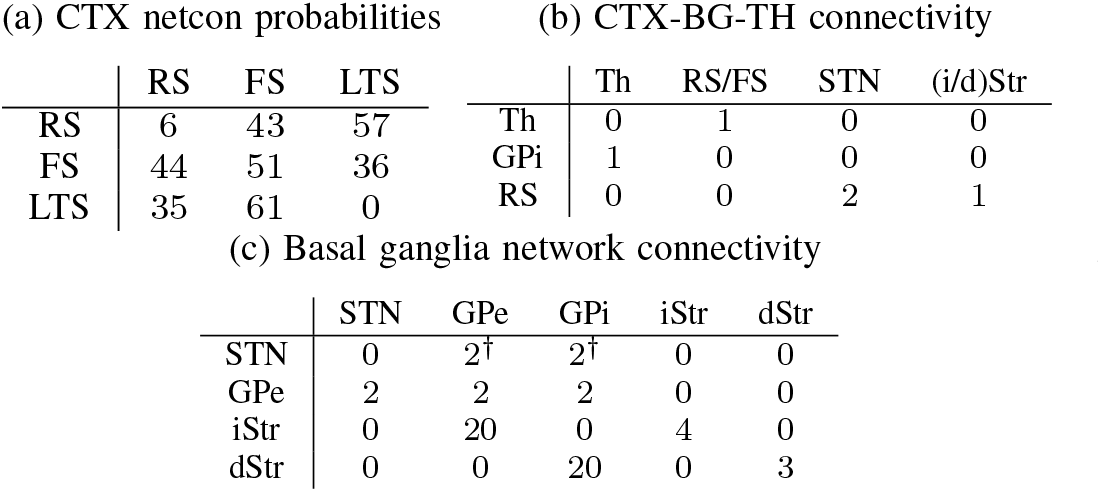
Cortex-Basal Ganglia-Thalamus Network topology. Source and target neurons respectively listed along rows and columns. (a) Probabilities in %. (b-c) Number of connections. (†) 50% of the target cells do not receive afferent input.

#### 2) Synaptic modeling

The synaptic current *I*_syn_ is given by the product of the maximal synaptic gain *g*_syn_, a synaptic parameter *s*, a plasticity-factor *P*, and a potential term *V-E*_syn_, with *E*_syn_ the GABAergic or glutamatergic nernst potential:

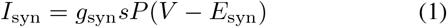

Synaptic gains *g*_syn_ are summarized in Table II(a-c) for all connections in the Parkinsonian network.

**TABLE II:**
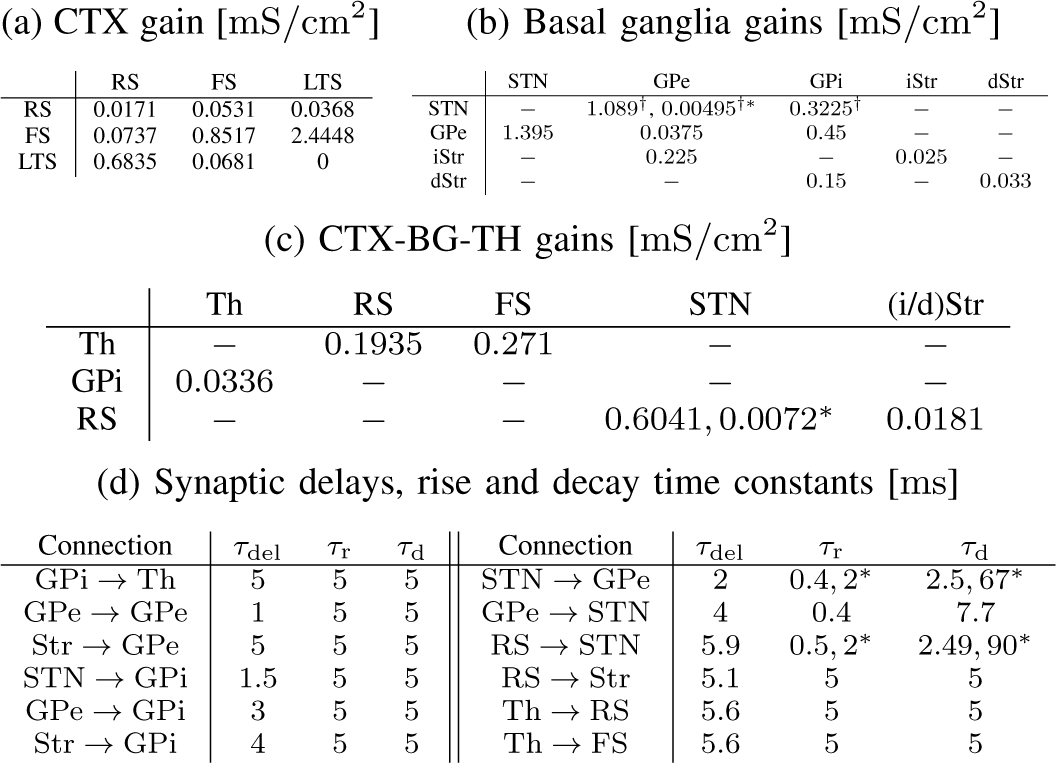
Synaptic delays and gains of the Parkinsonian neuronal network. Source and target neurons respectively listed along rows and columns. (a) Intracortical, (b) basal ganglia, and (c) cortex-basal ganglia-thalamus synaptical gains [mS/cm^2^] and (d) delays, rise and decay times [ms]. † Uniformly distributed around mean given in table * Gain or time constant of AMPA and NMDA, respectively.

The synaptic channel parameter s is activated by presynaptic spikes. For recurrent striatal GABA-A synapses, s is modeled by [36]:

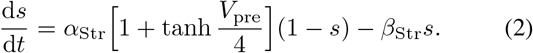

Here, *V*_pre_ is the presynaptic potential and *α*_str_ = 2 ms^-1^, *β*_Str_ = 0.0769 ms^-1^.

Cortical synapses are modeled as summed bi-exponentials [46]:

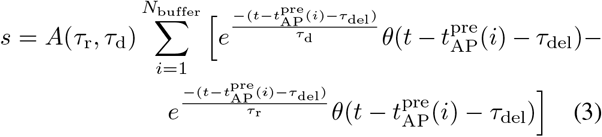

Here, *τ*_r_ and *τ*_d_ are the rise and decay time, respectively. *θ* is the Heaviside function and 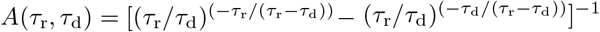 is a normalisation factor. Synaptic delays *τ*_del_ are summarized in Table II(d). Presynaptic action potential times are stored in a first in, first out buffer with size 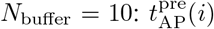 is the *i*th presynaptic spiking time in the buffer (spikes are detected by crossing of the threshold potential *V*_th_ = −10 mV).

Similarly, an alpha-synaptic current is a special case of a biexponential synapse (*τ*_d_ → *τ*_r_ in (3)) and is given by (the normalisation factor *A* reduces to the number of Euler *e*):

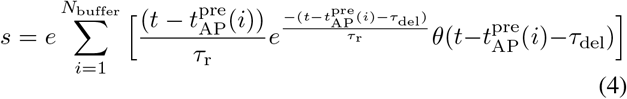

Glutamate synapses from pyramidal regular spiking (RS) neurons are AMPAergic (alpha-amino-3-hydroxy-5-methyl-4-isoxazolepropionic acid) and have rise and decay times of 0.1 ms and 3 ms, respectively. GABA-A (gammaaminobutyric acid) synapses from fast spiking (FS) interneurons have rise and decay constants of 0.5 ms and 8 ms, respectively. The GABA-A synapse from low threshold spiking (LTS) interneurons has a rise and decay constant equal to 0.5 ms and 50 ms [25], [46]. A synaptic delay of *τ*_del_ = 1 ms is introduced for cortical connections.

Short-term synaptic-plasticity is modeled as in [55], i.e., synaptic gains are multiplied with a plasticity-factor *P* = *F* (RS → LTS: facilitating connection) or *P* = *F* · *D*_1_ · *D*_2_ (RS → FS: depressing connection), cfr. Fig. 1 (*P* = 1 in connections without plasticity). Here, the dynamics of the facilitating factor *F* is given by:

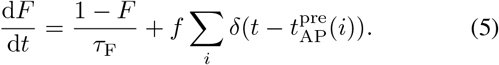

The sum is over the presynaptic spike times 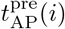 (*δ* is the dirac-delta distribution). Consequently, *F* will decay exponentially with time constant *τ*_F_ to one in the absence of presynaptic spikes (*τ*_F_ = 200 ms and *τ*_F_ = 94 ms for the RS → LTS and RS → FS connections, respectively). Conversely, after a presynaptic spike F is incremented with *f*(*f* = 0.2 and *f* = 0.5 for RS → LTS and RS → FS, respectively).

Similar dynamics holds for the depressing factor *D*, but here the effect of a presynaptic spike is multiplicative:

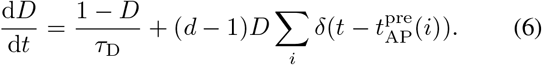

The time constants are *τ*_D1_ = 380 ms and *τ*_D2_ = 9200 ms. The multiplicative d-factors are equal to *d*_1_ = 0.46 and *d*_2_ = 0.975.

Finally, all glumatergic synapses have Nernst-potential *E*_AMPA_ = *E*_NMDA_ = 0 mV. Glutamate-synapses are AMPAergic, except for the STN → GPe connection and the hyperdirect pathway (RS → STN) that also contain NMDA-synapses. The GABAergic interstriatal connections have *E*_GABA,Str_ = −80 mV, while GABAergic Nernst-potentials in other nuclei are set to *E*_GABA_ = −85 mV. Synaptic delays, time constants and gains of the Parkinsonian network are summarized in Table II.

#### 3) Neuronal models

A modified Hodgkin-Huxley equation is used for the simulation of the neuronal response of the different nuclei:

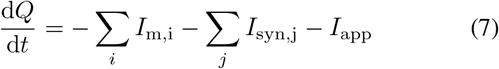

Here, *I*_m,i_; and *I*_app_ are the neuron-type specific membrane currents and externally applied current, respectively. Table III lists an overview of the membrane currents present in the different models and references to their dynamics. The current *I*_syn,j_ represents the received synaptic current from source population *j*.

**TABLE III:**
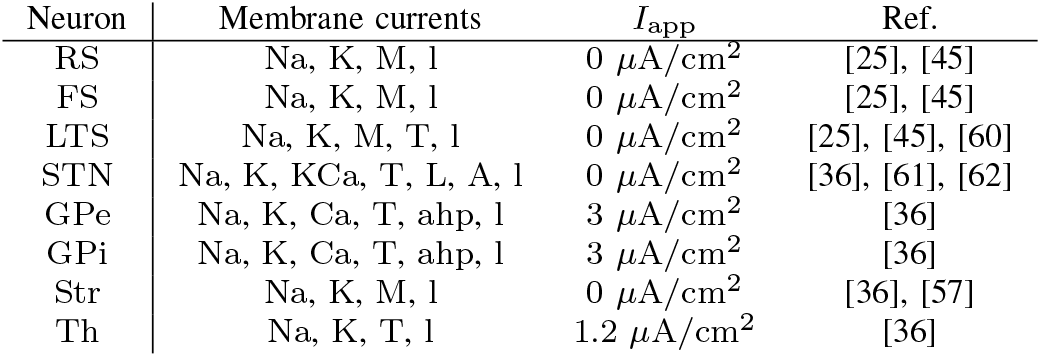
Summary of Neuron Models. Na: sodium; K: delayed-rectifier potassium; M: M-type slow noninactivating potassium; KCa: calcium dependent potassium; T: low-threshold T-type calcium; L: high-threshold L-type calcium; A: A-type potassium; Ca: high-threshold calcium; ahp: afterhyperpolarization; l: leak

| Neuron | Membrane currents | $I_{app}$ | Ref. |
| --- | --- | --- | --- |
| RS | Na, K, M, l | $0 \mu\text{A}/\text{cm}^2$ | [25], [45] |
| FS | Na, K, M, l | $0 \mu\text{A}/\text{cm}^2$ | [25], [45] |
| LTS | Na, K, M, T, l | $0 \mu\text{A}/\text{cm}^2$ | [25], [45], [60] |
| STN | Na, K, KCa, T, L, A, l | $0 \mu\text{A}/\text{cm}^2$ | [36], [61], [62] |
| GPe | Na, K, Ca, T, ahp, l | $3 \mu\text{A}/\text{cm}^2$ | [36] |
| GPI | Na, K, Ca, T, ahp, l | $3 \mu\text{A}/\text{cm}^2$ | [36] |
| Str | Na, K, M, l | $0 \mu\text{A}/\text{cm}^2$ | [36], [57] |
| Th | Na, K, T, l | $1.2 \mu\text{A}/\text{cm}^2$ | [36] |

#### 4) Parkinson’s disease and deep brain stimulation

The neuronal network of the cortex-basal ganglia-thalamus loop was made Parkinsonian, by three modifications to the healthy network. First, as in Kumaravelu et al. (2016) [36], the maximal M-type potassium current gain is decreased from *g*_M,H_ = 2.6 *μ*S/cm^2^ in the healthy (H) network to *g*_M,PD_ = 1.5 *μ*S/cm^2^ in the Parkinsonian (PD) condition, caused by heightened striatal acetylcholine concentrations after dopamine depletion [36], [56], [57]. Second, an additional external current of −2 *μ*A/cm^2^ is applied to the subthalamic nucleus and globus pallidus (current additional to *I*_app_ in Table III). Third, synaptic gains in the reciprocally coupled STN – GPe feedback loop and in the long loop (hyperdirect pathway and back from thalamus to cortex) are increased to promote propagation of beta oscillatory power into the Parkinsonian basal ganglia network. A similar methodology of altering synaptic gains and external applied currents to render the network Parkinsonian, has been used in other computational studies of the basal ganglia [34]–[38], [40], [54], [58]

Electrical deep brain stimulation is simulated by injection of a 300 *μ*A/cm^2^ current in the subthalamic nucleus cells with pulse duration 300 *μ*s and pulse repetition frequency between 10 Hz and 160 Hz [34]–[36], except when mentioned otherwise in the figure caption. Ultrasonic subthalamic nucleus deep brain stimulation is modeled with the multiScale Optimized model of Neuronal Intramembrane cavitation (SONIC) [59], cfr. section II-B.

#### 5) Spectral analysis and beta-oscillations

Parkinsonian bradykinesia is related with spectral density in the beta-band [63]–[66]. Here, we follow the hypothesis that cortical beta-band activity will enter the dopamine-depleted basal ganglia, where it is strenghtened by the STN-GPe feedback loop [37], [58], [64], [67]. Consequently, a beta-rhythm is imposed on the pyramidal regular spiking cortical neurons, by intracellular injection of 1 *μ*A/cm^2^ with normally distributed pulse repetition frequency (mean 20 Hz, standard deviation 3 Hz) and normally distributed duty cycle (mean 50%, standard deviation 25%).

Spectral analysis was performed by the multi-taper method (5 tapers, timebandwidth product equal to 3) in the chronux-Matlab toolbox (chronux.org, [68]). As in [36], spectograms are calculated with a sliding window of 1 s with step size 0.1 s and the alpha-beta energy is the integrated spectral density over the alpha-beta band (7 Hz – 35 Hz).

### B. Ultrasonic neuromodulation

Computational modeling of ultrasonic neuromodulation is based on the tentative bilayer sonophore (BLS) underlying mechanism (cfr. Fig. 1(b)) [23]. Here, the ultrasonic pressure wave is assumed to induce a sinusoidal oscillation of the deflection of both bilipid layer leaflets. Consequently, the membrane capacitance fluctuates in phase with the ultrasonic pressure wave, resulting in capacitive displacement currents. In the Neuronal Intramembrane Cavitation Excitation (NICE) model, these capacitive currents will result in membrane charge build-up, causing neuronal excitation [24], [25]. In this study, we adopt the NICE-framework for simulation of ultrasonic neuromodulation in isolated point neurons, as also used in previous studies [18], [23]–[25], [59], [69], [70]. The general model parameters are given in Table IV.

**TABLE IV:** Summary of UNMOD Parameters

| Abbr. | Value | Description |
| --- | --- | --- |
| $a_{BLS}$ | 32 nm | Radius bilayer sonophore |
| $C_{m0}$ | $1 \mu\text{F}/\text{cm}^2$ | Rest capacitance |
| $P_0$ | $10^5 \text{ Pa}$ | Static pressure |
| $T$ | $36 \text{ }^\circ\text{C}$ | Temperature |
| $\delta_0$ | 2 nm | Thickness leaflet |
| $\Delta$ | 1.26 nm | Intraleaflet gap at rest potential |
| $\mu_l$ | $0.7 \cdot 10^{-3} \text{ Pa} \cdot \text{s}$ | Dynamic viscosity of the CSF |
| $\mu_s$ | $0.035 \text{ Pa} \cdot \text{s}$ | Dynamic viscosity of leaflets |
| $\rho_l$ | $1028 \text{ kg}/\text{m}^3$ | Cerebrospinal fluid density |

The computational network model contains timescales across six orders of magnitude: a microsecond timescale of the ultrasound-neuron coupling, millisecond timescale of action potentials and membrane gating, and a second timescale of spike-frequency adaptation and network plasticity effects. Consequently, multi-scale optimization is required in order to prevent exorbitant simulation times. First, in the NICE-implementation [24], computational stiffness is decreased by the introduction of an update time step (here, *T*_up_ = 50 *μ*s) that decouples the mechanical (Rayleigh-Plesset) and electrodynamical (Hodgkin-Huxley) problems. Second, a multiScale Optimized model of Intramembrane Cavitation (SONIC) was proposed by [59], achieving significant further reductions of computational stiffness and simulation time. In the SONIC-model, effective voltage and rate parameters are pretabulated, the modified Hodgkin-Huxley equation is charge recasted and a Lennard-Jones fit is applied to the intramolecular pressure. The introduction of the SONIC-tables effectively removes the smallest microsecond timescale from the model, resulting in a three-order of magnitude reduction in simulation time, while maintaining qualitatively accurate results [59]. Consequently, the multi-scale optimized SONIC-model is used for all neuronal network simulations. For a description of the NICE and SONIC model, we refer to Plaksin et al. (2014, 2016) [24], [25] and Lemaire et al. (2019) [59], respectively.

## III. Results

### A. Interaction of ultrasonic and electrical neuromodulation in isolated neuron models

In this section, we investigate the potential benefits of simultaneous electrical stimulation and insonication in isolated cortical and subthalamic nucleus point neuron models. In Fig. 2, the interaction of ultrasound with electrostimulation is shown, by plotting the firing rate contours as function of the injected electrical current and ultrasonic intensity.

**Fig. 2:**
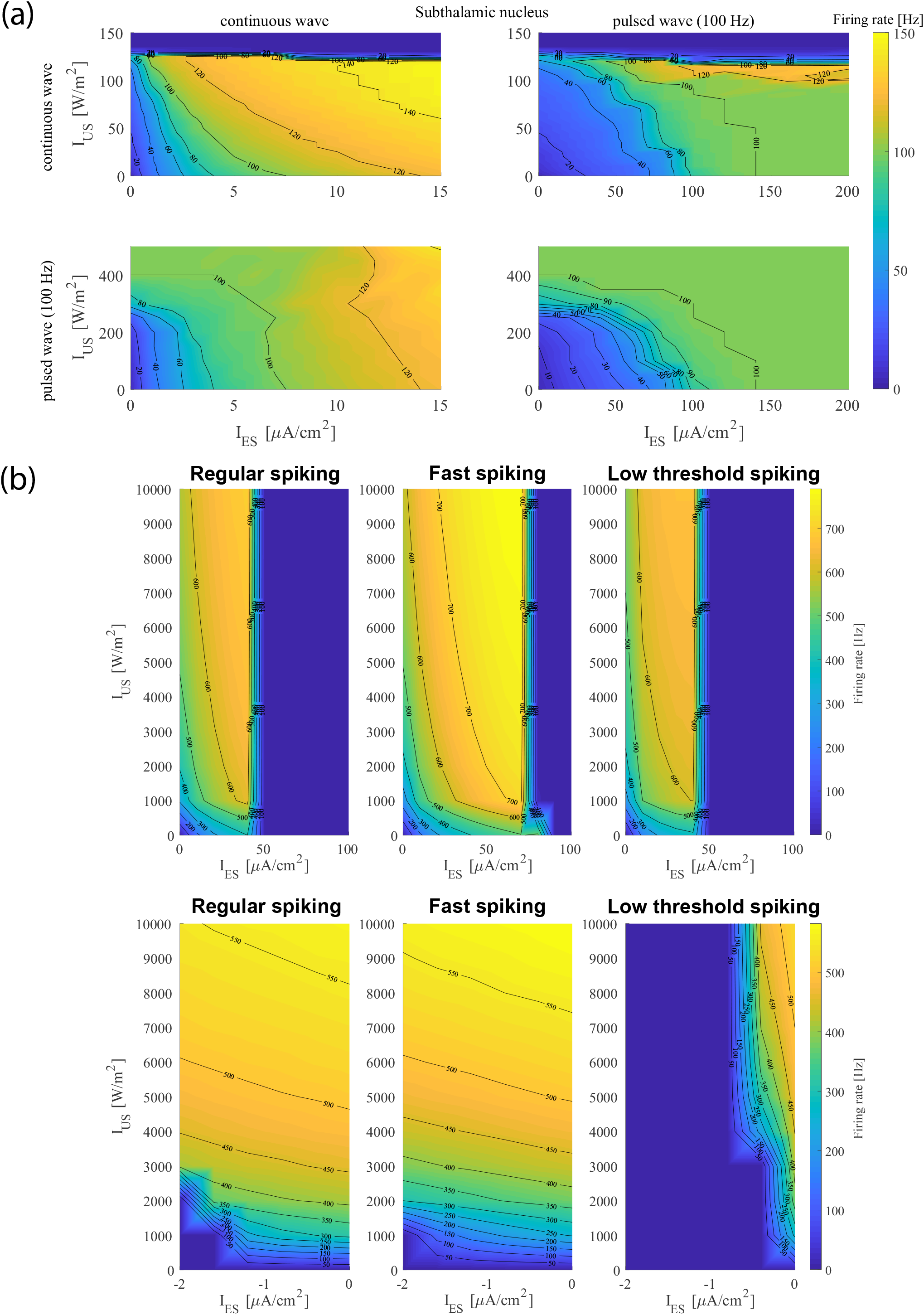
Combined ultrasonic and electrical neuromodulation firing rate contour plots. (a) Subthalamic nucleus response to simultaneous electrical and ultrasonic continuous wave or pulsed (PRF_US_ = PRF_ES_ = 100 Hz, f_US_ = 700 kHz) stimulation. Electrical and ultrasonic pulses are in phase (pulse durations *τ*_p,ES_ = 100 *μ*s, *τ*_P,US_ = 500 *μ*s). (b) Cortical regular spiking (left), fast spiking (middle) and low threshold spiking (right) response to simultaneous ultrasound and depolarizing (upper) and hyperpolarizing (lower) electrical currents

First, we investigate interacting ultrasound and electrical currents in a computational model of the plateau-potential generating subthalamic nucleus [61], [62] (Fig. 2(a)). A similar non-linear interaction between continuous-wave ultrasound and continuous (direct current) electrostimulation is observed for the STN (Fig. 2(a)(top,left)), compared to cortical cells (see below). However, here a silenced plateau is generated for 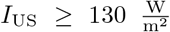, due to low-threshold T-type and high-threshold L-type calcium currents. This result is in agreement with our previous computational study on ultrasonic subthalamic nucleus stimulation [18]. Furthermore, combining pulsed ultrasound (PRF = 100 Hz) with DC electrical (Fig. 2(a)(bottom,left)) currents, or vice versa (Fig. 2(a)(top,right)), can achieve 100 Hz pulse-locked spiking at lower pulsed intensities/currents. However, increasing the DC-current or DC-intensity, will elicit neuronal spiking at rates higher than the pulse repetition frequency. In contrast, reliable pulse-locked spiking is achieved for simultaneous and in-phase pulsed electrical and ultrasonic neurostimulation (Fig. 2(a)(right,bottom)) at PRF = 100 Hz. Here, by simultaneous application of ultrasound with electrical current, 100 Hz spiking can be elicited with lower intensities or current injections.

Second, for depolarizing electrical current injections in cortical regular spiking, fast spiking and low-threshold spiking cells (Fig. 2(b)(upper)) a non-linear interaction is observed between ultrasound and the current injection. I.e., by combining ultrasound with electrostimulation, it is possible to maintain a given level of neuronal response (e.g., 400 Hz spiking) at a lower ultrasonic intensity and electrical current, compared to separately applied ultrasonic neuromodulation or electrostimulation. Here, we can also compare the neuronal response to ultrasonic insonication and electrostimulation applied separately: for electrical current injections the maximal spiking frequency is limited by depolarization block (region of silenced neuronal response at higher values of *I*_ES_ in Fig. 2(b)(upper)). In contrast, cortical neuronal response is not affected by depolarization block for the range of simulated ultrasonic intensities (up to *I*_US_ = 10 000 W/m^2^, which is above the FDA-limit on the temporal average peak intensity for diagnostic ultrasound (*I*_spta,FDA_ = 720 mW/cm^2^)) [71]. Consequently, the models predict that higher cortical firing rates are possible with ultrasonic neuromodulation than with electrostimulation.

Third, by combining ultrasound with hyperpolarizing electrical stimulation (Fig. 2(b)(lower)), it is possible to achieve a quiescent low threshold spiking cortical cell regime, while regular and fast spiking neurons exhibit high-frequency tonic spiking. In contrast, it is not possible to elicit high-frequency (e.g., 400 Hz) spiking of FS and RS-neurons without the excitation of LTS-cells, with non-simultaneous electrical currents or ultrasound. This region in Fig. 2(b)(lower) of silenced LTS-cells and spiking RS and FS neurons, obtained by combining ultrasonic and electrical neuromodulation, is explained by the observation that the LTS-neurons are most sensitive of the studied cortical cells to hyperpolarizing electrical currents. As a result, LTS-cells are inhibited faster than RS and FS neurons, while tonic spiking is maintained by insonication for the latter.

Finally, in Fig. 3(lower), firing rate contours of the subthalamic nucleus response to pulsed PRF = 100 Hz alternating electrical and ultrasonic stimulation are shown. For simultaneous application of alternating electrical depolarizing pulses (PRFES = 50 Hz) of sufficient amplitude to induce pulse-locked 50 Hz spiking and ultrasonic insonication with amplitudes at about 500 W/m^2^ (PRFUS = 50 Hz), neuronal spiking at 100 Hz is obtained. In Fig. 3 the different stimulation modalities (ultrasound and electrical currents) are separated in time (alternating pulses), in contrast with Fig. 2, where the concurrent electrical current and ultrasonic pressure wave stimulation modalities interact simultaneously. This separation of electrical and ultrasonic pulses is interesting from the perspective of safety, under the assumption that possible electrical/ultrasonic damage mechanisms are separated as well, and resolution (e.g., with similar ideas to intersectional short pulse stimulation [72]). From Fig. 3 it can also be observed that pulsed ultrasound alone at PRFUS = 50 Hz is not capable of reliable entrainment of neuronal spiking at 50 Hz. This observation agrees with our previous results [18], that pulse-locking for the subthalamic nucleus to ultrasound only occurs for PRF_US_ > PRF_min_ > 90 Hz.

**Fig. 3:**
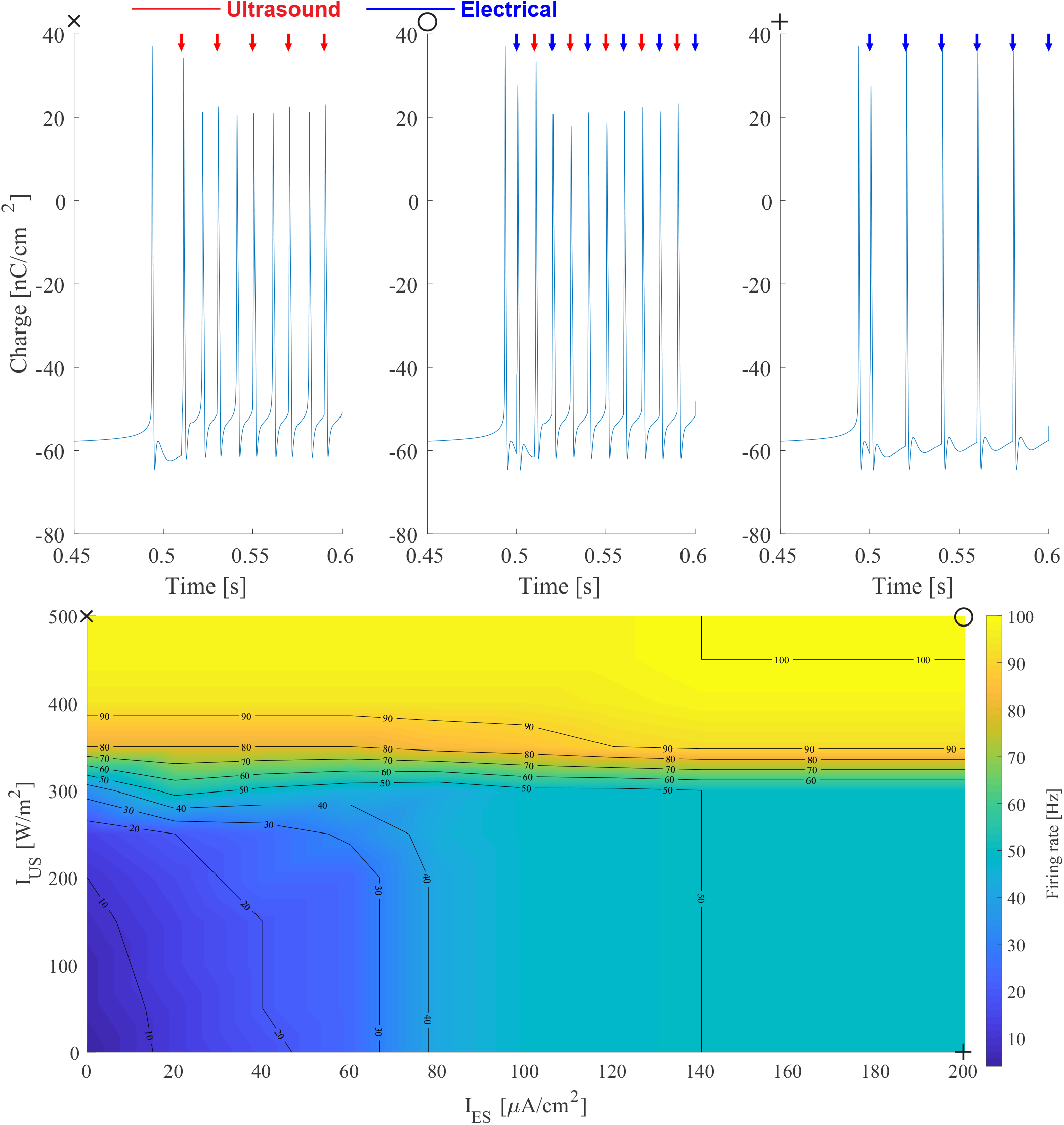
Subthalamic nucleus response to combined electrical and ultrasonic pulsed out-phase stimulation (pulse repetition frequency PRF_US_ = PRF_ES_ = 50 Hz, pulse duration *τ*_p,ES_ = 100 *μ*s, *τ*_p,US_ = 500 *μ*s, *f*_US_ = 700 kHz). (Upper) Membrane charge traces. Arrows indicate the timing of ultrasonic (red) or electrical (blue) pulses. (Lower) Combined ultrasonic and electrical neuromodulation firing rate contours. Markers indicate the waveform parameters for the corresponding membrane charge traces.

### B. Electrical and ultrasonic deep brain stimulation in the cortex-basal ganglia-thalamus computational network

#### 1) Ultrasonic and electrical stimulation separately

First, we compare the effects of ultrasonic and electrical subthalamic nucleus deep brain stimulation on the cortex-basal ganglia-thalamus network, when they are applied separately. Example rastergrams of the Parkinsonian network in the absence of neurostimulation and in the presence of electrical 160 Hz and ultrasonic deep brain stimulation are shown in Fig. 4. In Fig. 5(a-b,d-e,g-h), the dependency of the alpha-beta spectral energy (*α-β* SE), mean and standard deviation of the firing rate (*μ*FR and *σ*FR, respectively) are shown as function of the pulse repetition frequency in pulsed electrical and ultrasonic DBS (Fig. 5(a,d,g)) and as function of the ultrasonic intensity *I*_US_ for continuous-wave ultrasonic deep brain stimulation (Fig. 5(b,e,h)). In the Parkinsonian network, cortically imposed synchrony and beta-oscillations (e.g., see the RS trace in the PD rastergram in Fig. 4(left)) result in increased oscillations in the basal ganglia nuclei (cfr. Fig. 4(left), the parkinsonian spectrograms in Fig. 5(i,l), and the power spectral density plots in Fig. 5(c,f)). In Fig. 4, electrical and ultrasonic subthalamic nucleus deep brain stimulation are applied between *t*_start_ = 3.5 s and *t*_end_ = 8.5 s. Electrical pulsed 160 Hz deep brain stimulation entrains the subthalamic nucleus neurons at a firing rate equal to the pulse repetition frequency, with negligible variation between different STN-neuron traces (Fig. 4(middle): STN-trace). Consequently, globus pallidus neurons (GPi and GPe) that receive STN-afferents manifest tonic spiking, while globus pallidus neurons that do not receive these afferents are silenced via the GPe → GPe lateral inhibitory connections and the GPe → GPi-pathway (Fig. 4(middle): GPi and GPe). Furthermore, thalamic cells receive topologically-structured GPi GABAergic input. Therefore, the ThRT-rastergram is an inversion of the GPi-rastergram (e.g., Fig. 4(middle): ThRT-trace). The observations from the example rastergrams are confirmed and placed in context by the dependency of firing rates on pulse repetition frequency in Fig. 5(d,g): subthalamic nucleus and globus pallidus mean firing rate increases with electrical DBS pulse repetition frequency (Fig. 5(d)). While STN firing rate eventually saturates to the PRF, resulting in a decrease of the variability in firing between STN-neurons with PRF (Fig. 5(d,g)), the σFR of globus pallidus neurons increases with PRF (Fig. 5(g)). The alpha-beta spectral energy in the basal ganglia nuclei decreases with pulse repetition frequency (Fig. 5(a)), while a local worsening of *α-β* oscillatory power is observed around PRF = 30 Hz. The effect of 160 Hz electrical DBS on the power spectral density and the GPi-spectogram, is shown in Fig. 5(c) and (j), respectively.

**Fig. 4:**
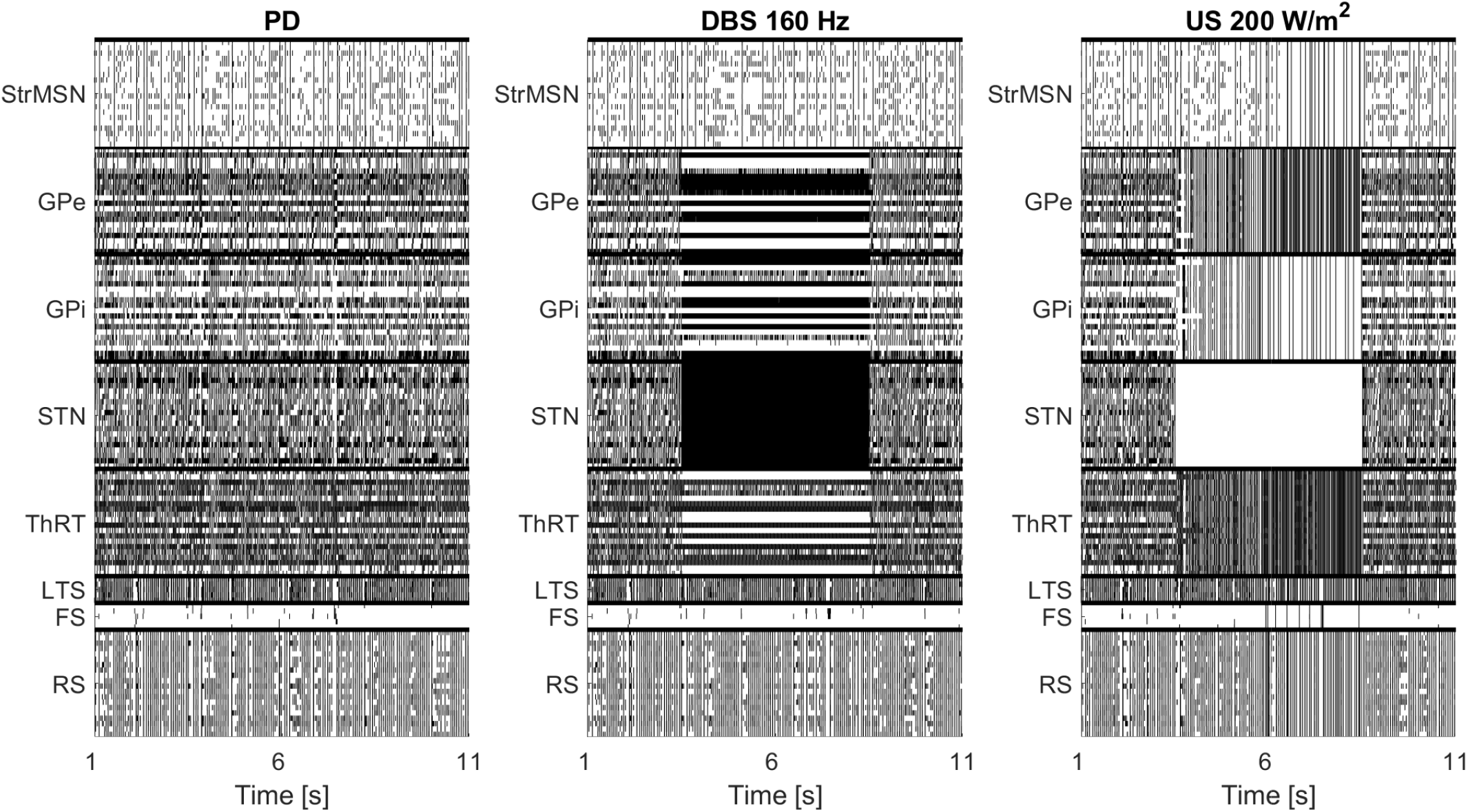
Action potential rastergram (vertical lines mark the timing of neuronal spikes) for the Parkinsonian (PD) cortex-basal ganglia-thalamus network without neuromodulation (left), with electrical deep brain stimulation (DBS: PRF_ES_ = 160 Hz, on time at 3.5 s, off time at 8.5 s; middle) and for continuous ultrasonic neuromodulation (US: 200 W/m^2^, *f*_US_ = 500 kHz, on time at 3.5 s, off time at 8.5 s; right). **Abbreviations.** StrMSN: striatal medium spiny neuron, GPe: globus pallidus external, GPi: globus pallidus internal, STN: subthalamic nucleus, ThRT: thalamic rubin-terman neuron, LTS: low threshold spiking cortical cell, FS: fast spiking cortical cell, RS: regular spiking cortical cell.

**Fig. 5:**
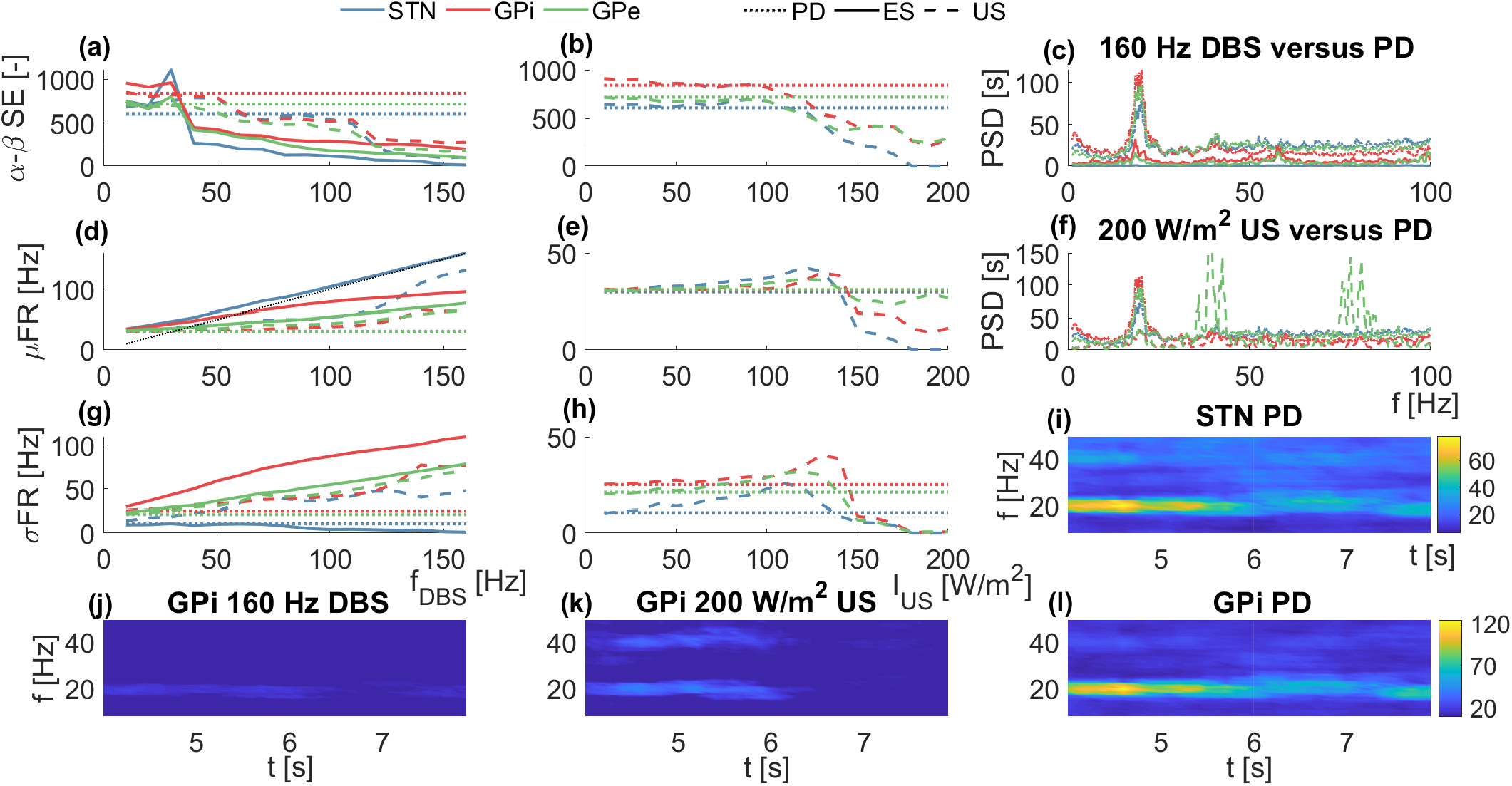
Electrical and ultrasonic (*f*_US_ = 500 kHz) pulsed and continuous subthalamic nucleus deep brain stimulation: ES and UNMOD are applied separately. (a-h) Parkinsonian network without neuromodulation represented by dotted lines. (a) Electrostimulation (full) or pulsed ultrasound (*τ*_p,US_ = 500 *μ*s, *I*_US_ = 400 W/m^2^) (dashed) frequency-dependent effect on alpha-beta spectral energy (*α – β* SE). (b) Continuous ultrasound (dashed) intensity-dependent impact on *α – β* SE. (c) Power spectral density (PSD) during 160 Hz electrical DBS. Mean (d-e) (*μ*FR) and standard deviation (g-h) σFR of the firing rate versus the pulse repetition frequency (d,g) and intensity in continuous-ultrasound (e,h). (f) PSD during 200 W/m^2^ continuous ultrasound. (i-l) Spectograms in the Parkinsonian subthalamic nucleus (STN) (i) and globus pallidus internus (GPi) (l) without neuromodulation. GPi spectogram during electrical 160 Hz DBS (j) and continuous UNMOD (k) ((j-l) are on the same colour scale).

The results for ultrasonic pulsed deep brain stimulation are similar to electrical pulsed DBS (Fig. 5(a,d,g), dashed). Alphabeta spectral energy desirably reduces with pulse repetition frequency: here, no local worsening at low pulse repetition frequencies is observed for the GPi and GPe spectral energies, while the increase in STN beta-oscillations at low PRF is less pronounced. Although STN firing increases with ultrasonic PRF, no complete entrainment of subthalamic nucleus neurons at the pulse repetition frequency is observed for *I*_US_ = 400 W/m^2^ (cfr. (Fig. 5(d)), dashed with dotted line). For PRF = 160 Hz and *I*_US_ = 400 W/m^2^, 25% of the STN-neurons have a firing rate that is not within PRF±10%PRF. The neurons that are not yet pulse-locked receive strongly varying GABAergic GPe input, while entrained neurons receive a more constant low or high level of afferent inhibitory GPe current. At higher ultrasonic intensities all neurons become pulse-locked, e.g., for *I*_US_ = 500 W/m^2^ and PRF = 160 Hz, 100% of the STN-neurons fire at 160± 1 Hz. Here, we want to illustrate in Fig. 5(a) that *α – β* spectral energy is already improved, without complete STN pulse-locking to the ultrasonic stimulus.

Our model results predict that continuous-wave ultrasound acts by silencing subthalamic nucleus neurons to a plateau (Fig. 4(right, STN trace), Fig. 5(e)(blue dashed)). The lack of STN activity results in highly synchronized GPi and GPe activity in our network model (Fig. 4(right), GPe and GPi trace and *σ*FR drops to zero at higher ultrasonic pressures in Fig. 5(h)). However, alpha-beta spectral energy and firing rates are still reduced in the STN, GPi and GPe (Fig. 5(h) and Fig. 5(e), respectively). Instead, globus pallidus external oscillatory power is increased outside the alpha and beta bands, around 39 Hz and 78 Hz (Fig. 5(f), green dashed). The effect of continuous-wave ultrasound on the parkinsonian GPi-spectogram is shown in Fig. 5(gk).

#### 2) Ultrasonic and electrical stimulation in tandem

The application of simultaneous electrical and ultrasonic neuromodulation for the suppression of *α – β* spectral energy in the Parkinsonian network is presented in Fig. 6 and Fig. 7 for maximally out-of-phase and in-phase pulsed waveforms, respectively. From Fig. 6, we observe that STN, GPi and GPe *α – β*-oscillations decrease with the temporally alternating electro-acoustic stimulation frequency *f*_EUS_, i.e., the pulse repetition frequency of the outphase electrical or ultrasonic pulse trains (*f*_US_ = 500 kHz, *τ*_p,US_ = 500 *μ*s, *τ*_p,ES_ = 300 *μ*s, *I*_US_ = 500 W/m^2^, *I*_ES_ = 300 *μ*A/cm^2^). In line with separated pulsed ES and US (Fig. 5(d),(g)), mean firing rates increase with frequency (Fig. 6(b)). Here, the firing rate variability decreases and increases with pulse repetition frequency, for the subthalamic nucleus and the globus pallidus, respectively (Fig. 6(c)). The trends for the spectral energy, mean and standard deviation of the firing rate for alternating electrical and ultrasonic pulses with fEUS are in line with these for ultrasound or electrostimulation applied separately, but with twice the PRF (i.e., *f*_DBS_ = 2*f*_EUS_, indicated with the upper x-axis in Fig. 6(a-c)).

**Fig. 6:**
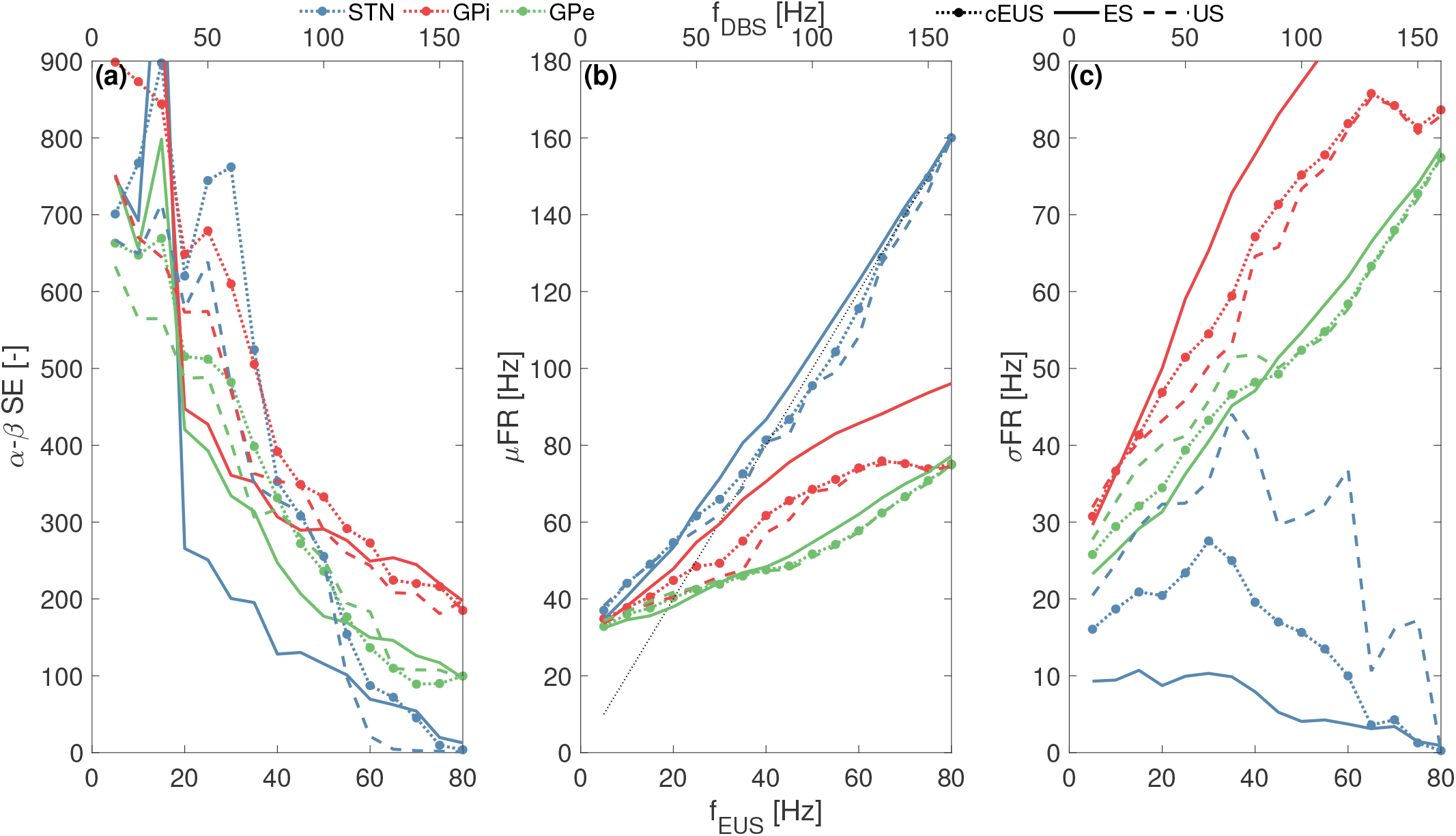
Effect of temporally alternating electrostimulation and ultrasonic (500 kHz, *τ*_p,US_ = 500 *μ*s) neuromodulation (cEUS, dotted, with marker) on alpha-beta spectral energy (a), mean firing rate (μFR) (b) and standard deviation of the firing rate (σFR) (c), in comparison with electrostimulation (ES, full lines) and insonication (US, dashed lines) applied separately.

**Fig. 7:**
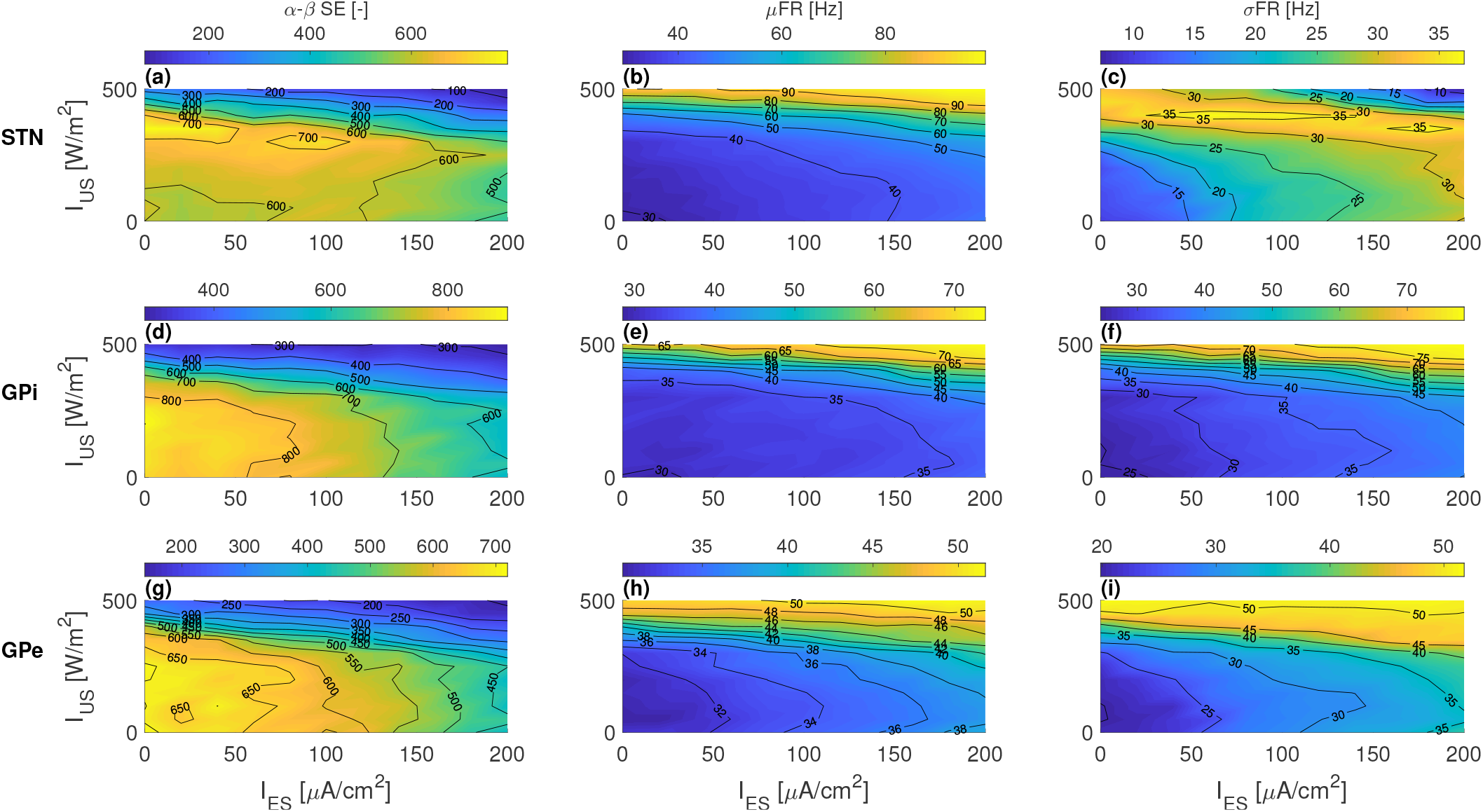
Effect of phase-locked electrostimulation (*τ*_p,ES_ = 100 *μ*s, PRF_ES_ = 100 Hz) and ultrasonic insonication (700 kHz, *τ*_p,US_ = 500 *μ*s, PRF_US_ = 100 Hz) on alpha-beta spectral energy (a,d,g), mean firing rate (*μ*FR) (b,e,h) and standard deviation of the firing rate (*σ*FR) (c,f,i) as function of the electric current amplitude and ultrasound intensity in the subthalamic nucleus (a-c), globus pallidus interna (d-f) and globus pallidus externa (g-i).

Next, in Fig. 7 phase-locked electro-ultrasonic stimulation is applied to the subthalamic nucleus neurons in the network with waveform parameters corresponding to Fig. 2(a,bottom right) (i.e., PRF_ES_ = PRF_US_ = 100 Hz, fuS = 700 kHz, *τ*_p,ES_ = 100 *μ*s and *τ*_p,US_ = 500 *μ*s). Here, it is interesting to observe the discrepancy in the firing rate contours of the subthalamic nucleus (cfr. Fig. 2(a,bottom right) and Fig. 7(b)). In particular, while in the isolated STN 100 Hz firing was observed in a significant area of the explored parameter space (e.g., for *I*_ES_ ≥ 150 *μ*A/cm^2^, Fig. 2), similar pulse-locked firing (100 Hz ± 1%) is only reached at two simulation points ((*I*_ES_,*I*_US_) = (180 *μ*A/cm^2^, 500 W/m^2^), (200 *μ*A/cm^2^,500 W/m^2^)) for the mean firing rate of STN-neurons that receive synaptic afferents. Furthermore, the variation in firing rate σFR increases with electrical and ultrasonic strength in the parameter regions where pulse-locked mean firing is not achieved (Fig. 7(c)), indicating that the GABAergic and glutamatergic synaptic currents in the network model result in a reduction in reliable spiking and pulse-locking, compared to isolated STN-neurons. The GPi (Fig. 7(e,f)) and GPe (Fig. 7(h,i)) mean and standard deviation on the firing rate increase with the interacting electrical and ultrasonic intensity. Finally, we can observe from Fig. 7(a,d,g) that the Parkinsonian *α – β* oscillations are reduced faster by combining electrical and ultrasonic stimulation (i.e., spectral energy contours are not horizontal or vertical).

## IV. Discussion

### A. Combined UNMOD and electrostimulation

First, interaction of simultaneous (Fig. 2) and nonsimultaneous (Fig. 3) ultrasound and electrical currents was investigated in isolated neuron models. We observe that nonlinear interaction of UNMOD with electrostimulation can achieve a given level of neuronal response (firing rate contour), at lower ultrasonic intensity and electrical current than would be required if these stimulation modalities are used separately. These results predict an increase in firing rate dynamic range for EUS-stimulation, or conversely an improvement of safety, with respect to damage mechanisms that are separated for ultrasound and electrostimulation. For example, the probability of damage by ultrasound-induced inertial cavitation and via electrochemical effects is likely not influenced by the presence of electrical current and ultrasound, respectively. Another potential application of the observed interaction of ultrasound with electrical currents, is the construction of stimulation modalities that leverage this interaction for improved targeting resolution (e.g., combined TI-DBS and focused ultrasound with partially or fully overlapping foci). In theory, any electrostimulation technology (transcranial magnetic stimulation (TMS), transcranial direct current stimulation (tDCS), electrical deep brain stimulation (ES-DBS), etc.) can be combined with transcranial focused ultrasound (tFUS) in order to combine the benefits and mitigate the downsides of the neurostimulation methods. E.g., tFUS has the benefit of high millimeter size resolution and transcranial focusing, while electrostimulation is energetically more favourable (ultrasound requires six orders of magnitude more energy than direct current injection for action potential initiation [24]). Furthermore, our simulations predict that it is possible to achieve altered relative spiking regimes, by combining continuous UNMOD with hyperpolarizing currents. Here, RS and FS cell high-frequency tonic spiking activity could be induced in the absence of LTS-cell activity with EUS, which was not possible for continuous ultrasound or electrostimulation separately. This observation could have applications for therapy or research, because low threshold-spiking cells are thought to protect the cortical network against overexcitation [47]. In other words, combining ultrasound with hyperpolarizing currents might be an efficient approach to induce a cortical seizure.

### B. Deep brain stimulation in a CTX-BG-TH neuronal network

In this study, a biophysically realistic computational model of the cortex-basal ganglia-thalamus loop was constructed, based on [25], [34]–[36], [46], [47], [54], and its interaction with ultrasound and electrostimulation was investigated. Firing rates in the healthy and Parkinsonian network are in corre-spondence with recordings in healthy and 6-OHDA dopamine depleted rats [73]. The Parkinsonian network demonstrated increased basal ganglia oscillatory beta power, that is reduced by deep brain stimulation to the STN. Here, deep brain stimulation efficiency improves with increasing pulse repetition frequency between 50 Hz and 130 Hz, as observed in patients [74] and other basal ganglia models [35], [36]. Similar to electrical DBS, ultrasonic pulsed stimulation improves pathological network oscillations by driving the subthalamic nucleus spiking rate (Fig. 4–5(a,d,g)). Furthermore, interaction of temporally alternating or simultaneous ultrasonic and electrical pulses results in a higher obtained firing rate or reduction of Parkinsonian oscillations for given separated limits on electrical currents and ultrasonic intensity (Fig. 6–7). Here, we observe that a dichotomous response of globus pallidus neurons is obtained at therapeutic parameters (Fig. 6(c), Fig. 7(f,i)), similar to Kumaravelu et al. (2016) [36]: i.e., some globus pallidus neurons exhibit tonic spiking, while others are silenced. This is not unexpected, since our basal ganglia network is based on [36], in particular the presence of two types of globus pallidus neurons, distinguished by whether or not they receive subthalamic afferents (cfr. Table I(c)).

Conversely, continuous-wave ultrasound also reduces alphabeta oscillations in our model, but by silencing the STN to a plateau-potential (Fig. 4–5(b,e,h)). Here, silencing of the STN is functionally equivalent to subthalamotomy (i.e., ablation of the STN), which is an alternative treatment option for Parkinson’s disease, e.g., for patients that are unsuitable for DBS-lead implantation due to access or medical reasons [75]. Computational modeling of Hahn and McIntyre (2010) also predicts a decrease in pathological globus pallidus bursting upon silencing of STN neurons [76]. However, in contrast with this result, simulations of So et al. (2012) indicate a worsening of thalamic relay error indices by removing STN local cells alone [35], speculating that lesioning of the pallidothalamic pathway is required for symptom improvement, while lesions restricted to the STN might result in dyskinesias or hemiballism. Our simulations seem to agree with Hahn and McIntyre, in the sense that beta-oscillatory power is reduced upon STN-silencing with ultrasound. However, globus pallidus synchrony is increased and strong oscillations are observed outside the alpha-beta band. Here, we hypothesize that the exact response of proxies of Parkinsonian pathology to subthalamotomy might be sensitive to the values of the synaptic gains, network topology and externally applied currents. In line with this thought, So et al. observed that GPe bursting was dependent on lateral globus pallidus inhibition (*g*_GPe→GPe_) [35].

### C. Strengths, limitations and future work

In this paper, a fully biophysical ion-channel based neuronal network of the cortex-basal ganglia-thalamus loop has been used to quantify for the first time the applicability of ultrasonic neuromodulation for the treatment of Parkinson’s disease with the alpha-beta spectral energy proxy. Furthermore, simulations in isolated neuron models and the neuronal network in this study show the benefits of simultaneous and alternating ultrasonic and electric neuromodulation.

Some important limitations and directions for future work should be taken into account. First, simulations in the cortex-basal ganglia-thalamus network and isolated neurons, are performed in single-compartment point neurons. Consequently, spatial effects are not taken into account and electrical deep brain stimulation is simulated by direct current injection as in earlier models [34]–[37]. Important spatial effects for electrical deep brain stimulation are antidromic propagation and cortical invasion via the hyperdirect pathway [39], [40], coactivation of fibers of passage [77] (e.g., the lenticular fasciculus (GPi → Th) passes dorsally to the subthalamic nucleus), decoupling of the somatic and axonal STN response [78], etc. Further research will also be necessary in order to determine the importance of spatial effects in ultrasonic neuromodulation (e.g., location of the excitation node and its dependence on the ultrasonic waveform). However, the construction of a multicompartmental morphologically realistic model of ultrasonic neuromodulation by intramembrane cavitation is complicated by the computational stiffness of the NICE-model, resulting in solver instabilities, low solution accuracy and/or exorbitant simulation times for neuronal models with a high number of compartments. For this reason, we designed a multi-scale optimized model SECONIC in [70] for the efficient integration of multi-compartmental UNMOD-BLS models, including fast charge oscillations and their impact on neuronal excitability. As future work, we intend to apply SECONIC to morphologically realistic models, allowing us to investigate computationally the spatial aspects of UNMOD. In this context, a recent computational study of Lemaire et al. (2020) investigated ultrasonic neuromodulation by intramembrane cavitation in multi-compartmental myelinated and unmyelinated axons with the SONIC-framework [79]. Here, spatially-extended multicompartmental neuron models could be integrated within the point neuronal network (as done in [58], for a cortical multi-compartmental cell) and coupled with finite-element and finite-difference time-domain simulations of the electric and ultrasonic field, respectively.

Second, the underlying mechanism for UNMOD is not well-understood. Here, we focused on the proposed bilayer sonophore mechanism [23], [24], in which oscillating intramembrane cavities result in capacitive displacement currents and neuronal excitation. However, several other tentative mechanisms have been proposed. E.g., instead of exerting its influence by the harmonic pressure component as in the BLS-mechanism, ultrasound could modulate neuronal activity via the acoustic radiation force [26], [27]. The acoustic radiation force mechanism could also be mediated via its effects on the membrane capacitance [28]. Mechanosensitivity of ion channels is another important tentative mechanism [29], [30], [80], in which the harmonic pressure component or acoustic radiation force alters the dynamics of the ion channels. Here, interaction with the bilayer sonophore model or Prieto-model of acoustic radiation force effect is possible: e.g., the NICE-model predicts oscillating membrane tension, which could impact the mechanosensitive ion channels. Furthermore, also flexoelectricity [31], thermodynamic neuron models [33], mechanical surface and axial waves [81], [82], propagation via acto-myosin cytoskeletal elements [83], etc., could be important for a complete understanding of ultrasonic neuromodulation. Here, computational modeling can be helpful to improve understanding of the consequences and potential interactions of the different tentative mechanisms. In this study, predictions have been made on the interaction of bilayer sonophore mediated ultrasonic neuromodulation and electrical currents, the comparative efficiency for the reduction of beta-oscillations of UNMOD w.r.t. electrostimulation, efficacy of continuous-wave ultrasound versus pulsed ultrasound, etc. These predictions could be tested in future experimental studies. As future work, we intend to incorporate also other tentative mechanisms of ultrasound (acoustic radiation force, mechanosensitivity of ion channels…) into the models, in order to investigate the interactions between them and to gain understanding under which conditions (ultrasonic waveform, neuron type,…) a certain mechanism is expected to be dominant.

## V. Conclusion

In this study, a computational biophysical (i.e., fully Hodgkin-Huxley) neuronal network of the Parkinsonian cortex, basal ganglia and thalamus was constructed and coupled with ultrasound and electrostimulation. Here, the basal ganglia receive cortical input via the hyperdirect and (in)direct pathways, while the thalamus relays the output from the globus pallidus back to regular spiking pyramidal cortical cells and fast spiking interneurons. Both electrical and ultrasonic pulsed STN-stimulation is capable of reducing elevated alpha-beta pathological oscillations at higher pulse repetition frequencies. Continuous-wave ultrasound is also able of improving alphabeta power, but by the opposing mechanism of silencing the subthalamic nucleus to an elevated plateau-potential. In both the neuronal network and in isolated neuron models it was observed that simultaneous or alternating application of electro-acoustic waveforms is capable of obtaining higher firing rates and better reduction in Parkinsonian oscillations for given safety limits on the electrical current and ultrasonic intensity. Furthermore, it was demonstrated in cortical cells that combining hyperpolarizing (negative) electrode currents with ultrasound can achieve altered relative cortical spiking regimes: i.e., entraining regular and fast spiking neurons to a high-frequency pulse train, while low-threshold spiking cells are quiescent.

The results presented in this study indicate that the combination of electrical and ultrasonic modalities has potential to improve safety and dynamic range, targeting resolution, recruitment selectivity, and energy efficiency. Furthermore, it is predicted that both continuous and pulsed transcranial focused ultrasound targeted to the STN is able to improve Parkinsonian symptoms. Our computational model provides novel testable predictions, that can be used to verify or falsify the proposed underlying mechanisms of DBS and ultrasound-neuron interaction.

As future work, we intend to perform a sensitivity study of the efficiency of ultrasonic and electrical stimulation to variations in the network topology and synaptic gains. Here, these variations could potentially be interpreted as inter-subject variability. Furthermore, target locations other than the STN (e.g., globus pallidus or thalamus) can be investigated, as well as co-stimulation of fibres of passage.

## VII. Acknowledgment

This work was carried out using the Supercomputer Infrastructure (STEVIN) at Ghent University, funded by Ghent University, the Flemish Supercomputer Center (VSC), the Hercules Foundation and the Flemish Government department EWI.

This research was funded by the FWO-project G046816N. T. Tarnaud is a PhD Fellow of the FWO-V (SB) (Research Foundation Flanders, Belgium). R. Schoeters is a PhD Fellow of the FWO-V (Research Foundation Flanders, Belgium).

